# Parametrising diffusion-taxis equations from animal movement trajectories using step selection analysis

**DOI:** 10.1101/2020.01.28.923052

**Authors:** Jonathan R. Potts, Ulrike E. Schlägel

## Abstract

1. Mathematical analysis of partial differential equations (PDEs) has led to many insights regarding the effect of organism movements on spatial population dynamics. However, their use has mainly been confined to the community of mathematical biologists, with less attention from statistical and empirical ecologists. We conjecture that this is principally due to the inherent difficulties in fitting PDEs to data.
2. To help remedy this situation, in the context of movement ecology, we show how the popular technique of step selection analysis (SSA) can be used to parametrise a class of PDEs, called *diffusion-taxis* models, from an animal’s trajectory. We examine the accuracy of our technique on simulated data, then demonstrate the utility of diffusion-taxis models in two ways. First, we derive the steady-state utilisation distribution in a closed analytic form. Second, we give a simple recipe for deriving spatial pattern formation properties that emerge from inferred movement-and-interaction processes: specifically, do those processes lead to heterogeneous spatial distributions and if so, do these distributions oscillate in perpetuity or eventually stabilise? The second question is demonstrated by application to data on concurrently-tracked bank voles (*Myodes glareolus*).
3. Our results show that SSA can accurately parametrise diffusion-taxis equations from location data, providing the frequency of the data is not too low. We show that the steady-state distribution of our diffusion-taxis model, where it exists, has an identical functional form to the utilisation distribution given by resource selection analysis (RSA), thus formally linking (fine scale) SSA with (broad scale) RSA. For the bank vole data, we show how our SSA-PDE approach can give predictions regarding the spatial aggregation and segregation of different individuals, which are difficult to predict purely by examining results of SSA.
4. Our methods give a user-friendly way in to the world of PDEs, via a well-used statistical technique, which should lead to tighter links between the findings of mathematical ecology and observations from empirical ecology. By providing a non-speculative link between observed movement behaviours and space use patterns on larger spatio-temporal scales, our findings will also aid integration of movement ecology into understanding spatial species distributions.

## 1 Introduction

Partial differential equations (PDEs) are a principle workhorse for mathematical biologists (Murray, 2003). Their strength lies in both their utility in describing a vast range of biological systems, and the existence of many mathematical techniques for analysing them. For example, the theory of travelling wave solutions has been used to understand spreading-speeds and spatial distributions of invasive species (Kot *et al.*, 1996; Petrovskii *et al.*, 2002; Lewis *et al.*, 2016). Likewise, linear pattern formation analysis has been used for understanding animal coat patterns (Turing, 1952; Murray, 1981; Nakamasu *et al.*, 2009), vegetation stripes in semi-arid environments (Klausmeier, 1999; Sherratt, 2005), spatial predator-prey dynamics (Baurmann *et al.*, 2007; Li *et al.*, 2013), and many more examples from ecology and beyond (Kondo & Miura, 2010). There is also a zoo of advanced techniques for analysing PDEs, such as asymptotic analysis, weakly non-linear analysis, energy functionals, calculus of variations, and so forth (Evans, 2010; Murray, 2012). These have been used less frequently in an ecological setting, but their application is not without precedent (Eftimie *et al.*, 2009; Tulumello *et al.*, 2014; Potts & Lewis, 2016a).

Here, we are specifically interested in using PDEs to model animal movement. In this context, PDEs are valuable for understanding how patterns of utilisation distribution (the distribution of an animal’s or population’s space use) emerge from underlying movement processes. PDEs have been successfully applied in this regard to phenomena such as territory and home range formation (Lewis & Moorcroft, 2006; Potts & Lewis, 2014), flocking and herding (Eftimie *et al.*, 2007), organism aggregations (Topaz *et al.*, 2006), and spatial predator-prey dynamics (Lewis & Murray, 1993). They have also been used to understand animal motion in response to fluid currents (Painter & Hillen, 2015), insect dispersal (Ovaskainen *et al.*, 2008), and search strategies (Giuggioli *et al.*, 2009). In all these examples, the models are assumed to operate on timescales over which death and reproduction have minimal effect. On such timescales, the emergent spatio-temporal patterns of animal distributions are determined solely by the movement decisions of animals as they navigate the landscape. These decisions may be influenced by relatively static aspects of the environment (e.g. Giuggioli *et al.* (2009); Painter & Hillen (2015)) or the presence of other animals (e.g. Eftimie *et al.* (2007); Topaz *et al.* (2006)) or a combination of the two (e.g. Moorcroft *et al.* (2006)).

Despite their broad use by applied mathematicians in general, and their great success in understanding the emergent properties of ecological systems in particular, PDEs have been much less-used in empirical or statistical ecology. This is perhaps due to the difficulties of parametrising them from data. One can, in principle, construct a likelihood function for a PDE model given the data. This has been done, for example, in mechanistic home range analysis studies (Moorcroft *et al.*, 2006; Lewis & Moorcroft, 2006) and to understand insect dispersal through patchy environments (Ovaskainen, 2004; Ovaskainen *et al.*, 2008). However, fitting the likelihood function requires numerically solving the PDE for many different parameter values (Ferguson *et al.*, 2016). Such numerics can be both time consuming and technically difficult, essentially constituting a research subfield in its own right (Johnson, 2012; Ames, 2014). This is especially true when there are multiple interacting populations, due to the inherent non-linearities in the resulting PDEs, and also when the datasets are very large, as is increasingly the case (Hays *et al.*, 2016).

To test the theoretical advancements of PDE research against empirical observations, it is thus necessary to develop quicker and technically simpler methods for parametrisation. Several such methods have been developed to this end. For example, homogenisation techniques have been recently developed to simplify numerical solutions of reaction-diffusion equations (a class of PDEs), by separating time-scales in a biologically-motivated way (Powell & Zimmermann, 2004; Garlick *et al.*, 2011). Hefley *et al.* (2017) combined these methods with Bayesian techniques to parametrise reaction-diffusion equations efficiently and accurately from data on animal locations and disease transmission. However, these techniques rely on there being a biologically meaningful way to separate spatio-temporal scales, which is system-dependent. Furthermore it still requires numerical solutions of PDEs (albeit simplified ones), with all the technical baggage they can engender.

Likewise, the technique of gradient matching can also be used for rapid inference of differential equation models (Xun *et al.*, 2013; Macdonald & Husmeier, 2015). However, whilst this method can speed-up inference considerably, applying it to a movement trajectory (as is our present concern) requires interpolating between the data-points to give a smooth utilisation distribution. Indeed, the accuracy of the inference can be highly dependent upon the choice of this smoothing (Ferguson, 2018). Therefore it is necessary, when applying gradient matching to a trajectory, to try various smoothing procedures, which can be time consuming. Then, only if the procedures give similar results can one be confident about the outcome. As a consequence, gradient matching is best suited to data where there are sufficiently many individual organisms that the utilisation distribution can be reliably estimated with high accuracy, e.g. when studying cell aggregations (Ferguson *et al.*, 2016). However, in many studies of vertebrate animals’ movements, only a limited number of individuals can be tracked. It would therefore be advantageous to find a simpler, robust method of inference for parametrising PDEs, tailored to such animal tracking data.

To fill this gap, we show here that the oft-used technique of step selection analysis (Fortin *et al.*, 2005; Forester *et al.*, 2009; Thurfjell *et al.*, 2014; Avgar *et al.*, 2016) can be used to parametrise a class of PDEs called *diffusion-taxis equations* from animal tracking data. This is a class of advection-diffusion equation (sometimes called convection-diffusion) where the advection is up or down a gradient of some physical quantity (e.g. a gradient of resources). Such PDEs can describe animal movement in relation to external factors (e.g. landscape features or con- or heterospecific individuals) and hence make them a suitable model for animal movement in many situations. Step selection analysis (SSA) is already very widely-used, being both fast and simple to implement. Indeed, implementation has recently become even simpler thanks to the release of the `amt` package in `R` (Signer *et al.*, 2019), so using our method does not require significant new technical understanding by practitioners.

The resulting diffusion-taxis equations consist of two terms: (i) the diffusion term, which denotes the tendency for the animal locations to spread through time, and (ii) the taxis term, which encodes drift tendencies in the animal’s movement. Both terms may, in principle, vary across space, in particular in response to external factors such as habitat features, resources, predators, or conspecific individuals. As such, once one gets past the use of mathematical notation that may not be a standard part of an ecologist’s background, this is a very intuitive way to think about animal movement (Ovaskainen, 2004).

In this work, we give a simple recipe for converting the output of SSA into parameters for a diffusion-taxis equation. We then show how to use systems of such equations to understand both quantitative and qualitative features of emergent space-use patterns. In particular, we demonstrate how to derive the steady-state utilisation distribution (UD) in certain cases. This UD can be written in a closed-form, analytic expression, obviating the need for time-consuming numerics (Signer *et al.*, 2017). It describes the long-term space use of animals (i.e. their home ranges) and, in contrast to the mere SSA-derived parameter values, can be used to make rigorous predictions about space-use (Moorcroft & Barnett, 2008; Potts & Lewis, 2014). We also show how to predict whether the UD of an individual animal or a population will either (i) tend to a uniform steady-state (animal spread homogeneously across the landscape), (ii) reach a steady state with aggregation or segregation patterns, or (iii) be in perpetual spatio-temporal flux, never reaching a steady state.

Knowing when these emergent spatial distributions may arise from movement processes is vital for understanding spatial distributions of individuals within a population and ultimately species distributions. Individuals often use non-diffusive movement mechanisms (e.g. spatially explicit selection of locations based on resources or presence of conspecifics) which scale up to different space-use patterns such as homogeneous mixing, spatial aggregation/segregation, or dynamic spatio-temporal patterns (Potts & Lewis, 2019). Such movement decisions and resulting patterns challenge the assumption of well-mixed populations in traditional population models. This also has implications for demography, for example via density dependence or carrying capacities (Morales *et al.*, 2010; Riotte-Lambert *et al.*, 2017; Spiegel *et al.*, 2017), as well as interspecific interactions in communitites such as competition (Macandza *et al.*, 2012; Vanak *et al.*, 2013). As such, we encourage increased research effort in examining the effects of movement mechanisms on spatial patterns. We propose the tools developed through this paper and Schlägel *et al.* (2019) as a means to aid such examination. Although the mathematical justification for the techniques given here requires some technical expertise, the recipes for implementing these techniques do not require advanced mathematical understanding (being SSA plus some straightforward post-processing), so will be usable for a wide range of ecologists.

## 2 Methods

### 2.1 From step selection to diffusion-taxis

Suppose an animal is known to be at location **x** at time *t*. Step selection analysis (SSA) parametrises a probability density function, *p*_*τ*_ (**z**|**x**, *t*), of the animal being at location **z** at time *t* + *τ*, where *τ* is a time-step that usually corresponds to the time between successive measurements of the animal’s location (Forester *et al.*, 2009). For our purposes, the functional form of *p*_*τ*_ (**z**|**x**, *t*) is as follows

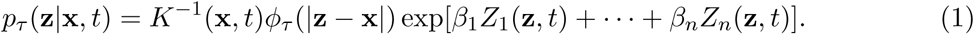

Here, *ϕ*_*τ*_ (|**z** − **x**|) is the step length distribution (i.e. a hypothesised distribution of distances that the animal travels in a time-step of length *τ*), |**z** − **x**| is the Euclidean distance between **z** and **x, Z**(**z**, *t*) = (*Z*_1_(**z**, *t*), …, *Z*_*n*_(**z**, *t*)) is a vector of spatial features that are hypothesised to co-vary with the animal’s choice of next location, ***β*** = (*β*_1_, …, *β*_*n*_) is a vector denoting the strength of the effect of each *Z*_*i*_(**z**, *t*) on movement, and

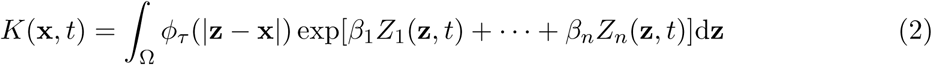

is a normalising function, ensuring *p*_*τ*_ (**z**|**x**, *t*) integrates to 1 (so is a genuine probability density function). In Equation (2), Ω is the study area, which we assume to be arbitrarily large. We also require that the step-length distribution, *ϕ*_*τ*_ (|**z** − **x**|), not be heavy-tailed (i.e. its mean, variance, and all its other moments must be finite). The parameters *β*_1_, … *β*_*n*_ are then the focus of an SSA, indicating the selection behaviour of animals towards spatial features of their environment. We refer to the function exp[*β*_1_*Z*_1_(**z**, *t*) + · · · + *β*_*n*_*Z*_*n*_(**z**, *t*)] as a step selection function (SSF), in line with its first use in the literature (Fortin *et al.*, 2005). Note, though, that sometimes SSF is instead used for the entire probability density function (Equation 1) (Forester *et al.*, 2009). SSA is the method of parametrising an SSF to analyse animal movement data.

One can generalise Equations (1-2) by incorporating environmental effects across the whole step from **x** to **z**, not just the end of the step at **z**. Furthermore, one can incorporate autocorrelation in movement via turning angle distributions (Forester *et al.*, 2009; Avgar *et al.*, 2016). However, for the purposes of parametrising an advection-diffusion PDE, both of these turn out to be unnecessary, so we stick with the functional form in Equation (1).

The SSA method requires data on a sequence of animal locations **x**_1_, …, **x**_*N*_ gathered at times *t*_1_, …, *t*_*N*_ respectively (with *t*_*j*+1_ − *t*_*j*_ = *τ* for all *j*, so that the time-step is constant), together with a vector of environmental layers, **Z**(**z**, *t*_*j*_) at each time-point *t*_*j*_. It then returns best-fit values for the parameters *β*_1_, …, *β*_*n*_, using a conditional logistic regression technique, by comparing each location with a set of ‘control’ locations sampled from an appropriate probability distribution, which represents locations that would be available to the animal based on its movement capabilities. Details of the SSA technique and how it should be implemented are given in previous works, e.g. Thurfjell *et al.* (2014); Avgar *et al.* (2016), so we omit them here.

We wish to use the SSA output to parametrise a diffusion-taxis model of the probability density function of animal locations, given by *u*(**x**, *t*). Notice that *u*(**x**, *t*) is different to the distribution described by Equation (1), which gives the probability density function of moving to location **z**, conditional on currently being located at **x**. However, in Supplementary Appendix B, we show that under the model in Equation (1), and as long as *τ* is sufficiently small, *u*(**x**, *t*) is well-described by the following diffusion-taxis equation

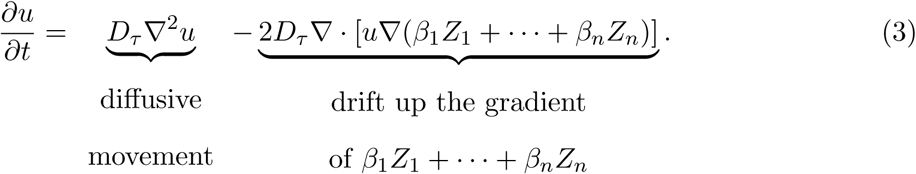

Here, ∇ = (*∂/∂x, ∂/∂y*) (where **x** = (*x, y*)), and

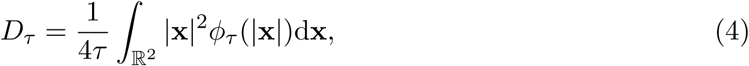

is a constant that describes the rate of diffusive movement. The derivation makes use of a diffusion-approximation approach (Turchin, 1998), whereby *u*(**x**, *t*) is derived by a moment-closure technique from a recurrence equation that describes how an animal’s location arises from its previous locations, and *p*(**z**|**x**) specifies the probability density of a specific movement step.

The drift part of Equation (3) describes animal movement in a preferred direction according to environmental features, whereas the diffusive part takes care of small-scale stochasticity due to any other factors not accounted for explicitly. For this approximation to work, *τ* must be sufficiently small that the gradient of resources (in any fixed direction) does not vary greatly across the spatial extent over which an animal is likely to move in time *τ* (see Supplementary Appendix B for precise mathematical details).

For our analysis, it is convenient to work in dimensionless co-ordinates. To this end, we start by setting 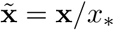 to be dimensionless space, where *x*_*_ is a characteristic spatial scale. Since, in practice, the functions *Z*_*i*_(**x**, *t*) arrive as rasterised layers (i.e. square lattices), it is convenient to let *x*_*_ be the pixel width (or, synonymously, the lattice spacing), but in principle the user can choose *x*_*_ arbitrarily. We also set 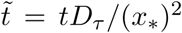 and 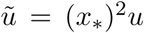. Then, immediately dropping the tildes above the letters for notational convenience, Equation (3) has the following dimensionless form

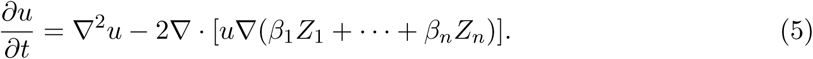

In summary, we have shown that step selection analysis can be used to parametrise a diffusion-taxis equation (Equation 5) where the drift term consists of taxis up the gradient of any covariate *Z*_*i*_ for which *β*_*i*_ is positive, and down the gradient of any covariate *Z*_*j*_ for which *β*_*j*_ is negative.

The key value in moving from the movement kernel in Equation (1) to the PDE in Equation (5) is that it allows us to make an explicit connection between a model, *p*_*τ*_ (**z**|**x**, *t*), of movement decisions over a small time interval, *τ*, and the predicted probability distribution, *u*(**x**, *t*), of an animal’s location at any point in time. While SSA by itself only gives inference about the movement rules themselves, the resulting PDEs enable us to make predictions of the space use patterns that will emerge over time, should the animal be moving according to the rules of the parametrised movement kernel. Examples of such patterns, including steady-state home ranges, aggregation, and segregation, will be demonstrated later in this manuscript.

### 2.2 Assessing inference accuracy on simulated data

To test the reliability of our parametrisation technique, we simulate paths given by diffusion-taxis equations of the general form in Equation (5). We then use step selection analysis to see whether the inferred ***β*** parameters match those that we used for simulations. For this study, we simulate two different types of model. In the first, which we call the *Fixed Resource Model*, there is just one landscape layer (so *n* = 1) and 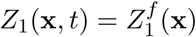 is a raster of resource values that does not vary over time (the superscript *f* emphasises that we are using the Fixed Resource Model). This raster is a Gaussian random field, constructed using the `RMGauss` function in the `RandomFields` package for `R`, with the parameter `scale=10` (Fig. 1a).

**Fig. 1.**
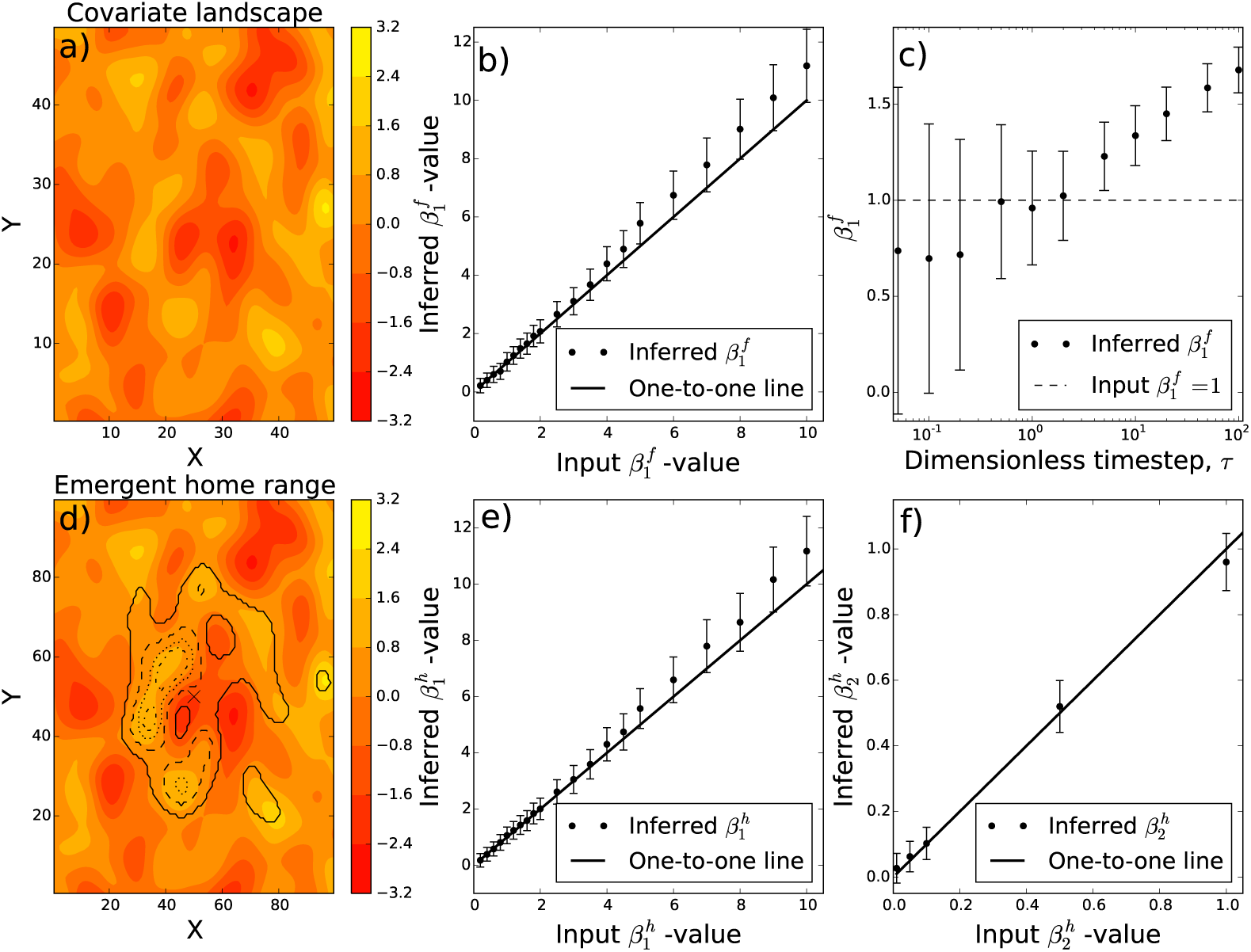
Study on simulated data. Inference from simulated paths of individuals moving according to the diffusion-taxis Equation (5). Panel (a) shows a resource layer given by a Gaussian random field, with colour showing the value of the resource layer at each point. Panel (b) gives the result of using step selection analysis to parametrise the Fixed Resource model, where 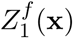 is given by this example layer. Dots give the inferred 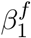 -values, with bars giving 95% confidence intervals. Panel (c) shows how inference varies as the time-step between measured locations, *τ* is increased. Here, the value used to simulate the diffusion-taxis equation is 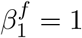. Panel (d) shows the emergent home range, as predicted by Equation (8), for the Home Range model with 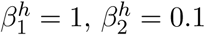. Here, 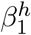 denotes the strength of the resource landscape’s effect on movement and 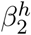 denotes the tendency to move towards the attraction centre, **x**_*c*_ (denoted by a cross). Details of this model are given in Section 2.2. The colour-filled contours are as in Panel (a) and the black curves show contours of the home range distribution. The solid black curve encloses 95% of the utilisation distribution. The 25%, 50%, and 75% kernels are given by dash-dot, dotted, and dashed curves respectively. Panels (e) and (f) show the results of using step selection analysis to infer 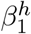 and 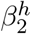, in an identical format to Panels (b) and (c). Panel (e) has 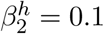 fixed and Panel (f) has 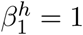 fixed.

The second model is called the *Home Range Model*. This has *n* = 2 (i.e. two landscape layers), the first of which, 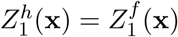, is the random field from Fig. 1a (the superscript *h* emphasises that we are working with the Home Range Model). The second denotes a tendency to move towards the central point on the landscape, which may be a den or nest site for the animal. This has the functional form 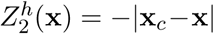, where **x**_*c*_ is the centre of the landscape. Notice that 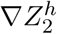 is an identical advection term to that in the classical Holgate-Okubo localising tendency model (Holgate, 1971; Lewis & Moorcroft, 2006).

For each of these two models, we simulate trajectories from Equation (5) for a variety of ***β***-values. Each trajectory consists of 1,000 locations, gathered at dimensionless time-intervals of *τ* = 1. (Recall from the non-dimensionalisation procedure that this corresponds to a time of 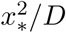 where *x*_*_ is the pixel width and *D* the diffusion constant of the animal, defined in Equation 4). We construct 10 trajectories for each ***β***-value used. Details of the method used for generating trajectories are given in Supplementary Appendix C. In short, the method involves reverse-engineering a stochastic individual-based model (IBM) from the PDE, such that the probability distribution of stochastic realisations of the IBM evolves in accordance with Equation (5). For the Fixed Resource Model, we also perform the same procedure but fixing 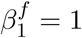 and varying *τ*, to understand the effect on inference of the time step, *τ*, at which data are gathered.

We then parametrise each trajectory using SSA, finding control locations by sampling steps from a bivariate normal distribution with zero mean and a standard deviation equal to the empirical standard deviation. We match each case to 100 controls. To determine whether SSA is effective in parametrising diffusion-taxis equations, we test whether the inferred ***β***-values fall within 95% confidence intervals of the values used to simulate the trajectories.

### 2.3 Application to empirical data and spatial pattern formation

To demonstrate the utility of diffusion-taxis models for animal movement, we used some recent results from a study of social interactions between bank voles (*Myodes glareolus*), reported by Schlägel *et al.* (2019). This study used SSA to infer the movement responses of each individual in a group to the other individuals. For example, individual 1 may tend to be attracted towards 2, who in turn may like to avoid 1 but rather be attracted towards 3. In the studied bank voles, such individualistic responses arose as sex-specific behaviours likely related to mating. However, they may also arise in relation to social foraging or interactions between species in competitive guilds.

Details of the method are given in Schlägel *et al.* (2019), but here we give the ideas pertinent to the present study. Suppose there are *M* individuals in a group. For each individual, *i*, consider the utilisation distribution of each of the other individuals to be a landscape layer. In other words *Z*_*j*_(**x**, *t*) = *u*_*j*_(**x**, *t*) in the step selection function (Equation 1). It may not be immediately obvious that one individual may be able to have knowledge about another’s utilisation distribution, but there are at least two biological processes by which this can happen, both of which can be justified mathematically (Potts & Lewis, 2019). The first is for individuals to mark the terrain as they move (e.g. using urine or faeces) and then the distribution of scent-marks mirrors the utilisation distribution (Gosling & Roberts, 2001; Potts & Lewis, 2016b). The second is for animals to remember past interactions with other individuals and respond to the cognitive map of these interactions (Fagan *et al.*, 2013; Potts & Lewis, 2016a).

By Equation (5), these movement processes give rise to a system of diffusion-taxis equations, one for each individual in the group, that each have the following form (in dimensionless coordinates)

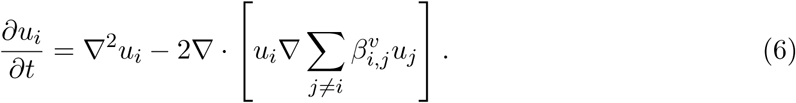

Here, 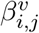 measures the tendency for individual *i* to move either towards (if 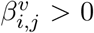) or away from (if 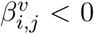) individual *j*. The magnitude of 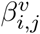 measures the strength of this advective tendency. These correspond to the ***β***-values inferred by SSA, with a superscript *v* to emphasise that these refer to the bank vole study.

Depending on the values of 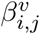, such a system of diffusion-taxis equations can have rather rich dynamics. These dynamics can be observed through numerical simulations (Fig. 2b). However, for technical reasons, to perform numerics we have to replace *u*_*j*_ in Equation (6) with a locally-averaged version 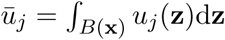, where *B*(**x**) is a small neighbourhood of **x**. This is to avoid rapid growth of small perturbations at arbitrarily high frequencies, which can happen without spatial averaging [see Supplementary Appendix E and Potts & Lewis (2019) for details]. The system we simulate is thus as follows

**Fig. 2.**
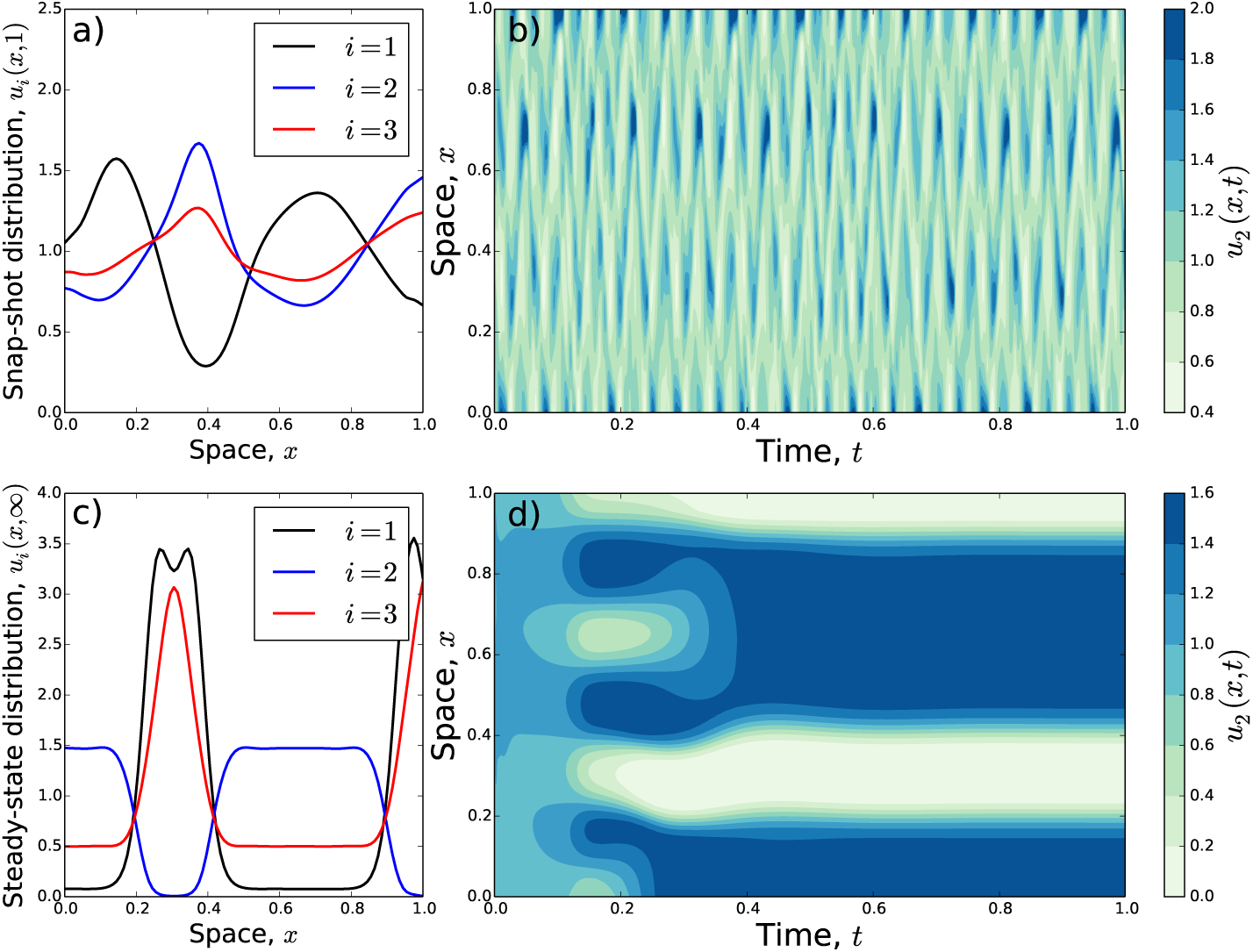
Pattern formation from diffusion-taxis systems. Panels (a) and (b) give a numerical solution of the system in Equation (7) in a simple one dimensional example, with *M* = 3 individuals (indexed with the letter *i*), 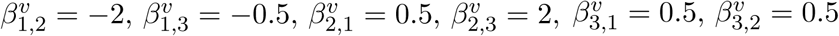. This is in the regime where linear pattern formation analysis predicts oscillatory patterns. Panel (a) gives a snap-shot of the system at *t* = 1, showing distributions of *u*_1_(*x*, 1), *u*_2_(*x*, 1), and *u*_3_(*x*, 1). Panel (b) shows the change in *u*_2_(*x, t*) over both space and time. We observe that the system never seems to settle to a steady state. This contrasts with Panels (c) and (d) which show a one dimensional example where linear pattern formation analysis predicts stationary patterns to emerge. Here, 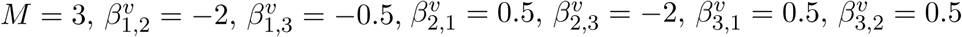. Panel (c) gives the stationary distribution, whilst Panel (d) displays convergence of the system towards this stationary distribution, for *u*_2_(*x, t*). Throughout all panels, the spatial averaging kernel is *B*(*x*) = (*x* − 0.05, *x* + 0.05) (see comment before Eqn. 7).

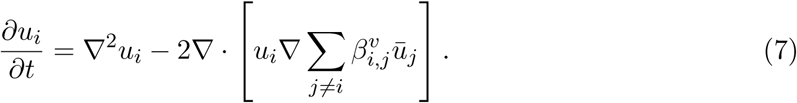

Details of the numerics are given in Supplementary Appendix E. To demonstrate some of the patterns that can emerge, Fig. 2 displays the spatio-temporal dynamics of the system in Equation (7) for various example parameter values. In Fig. 2a,b, we have *M* = 3, 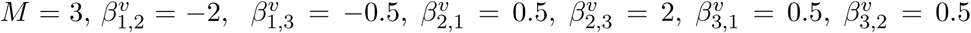. This means that Individual 1 is avoiding both 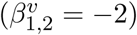 and 3 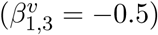; however 2 and 3 are both attracted towards 1 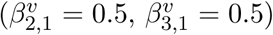 and also each other 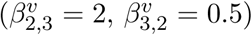. This complicated three-way relationship turns out to cause perpetually oscillating spatial patterns (Fig. 2a,b).

In Fig. 2c,d, we have *M* = 3, 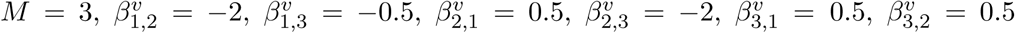. Thus Individual 1 still avoiding both 2 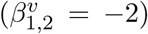 and 3 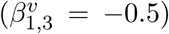. Furthermore, 2 and 3 are both still attracted towards 1 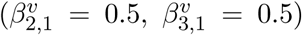 and 3 is attracted to 2 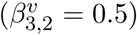. However, this time 2 is avoiding 3 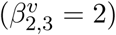. This situation leads to stationary spatial patterns (Fig. 2c,d).

It is perhaps not immediately obvious why this simple switch in behaviour from 2 being attracted to 3 to 2 avoiding 3 should have such a dramatic change in the qualitative nature of the utilisation distributions. However, one can gain insight into such effects by using linear pattern formation analysis (Turing, 1952). This technique separates parameter space into three regions: (a) *No Patterns*, so each individual will eventually use all parts of space with equal probability, (b) *Stationary Patterns*, where the system reaches a steady state such that individual utilisation distributions either become spatially segregated (with some possible overlap) and/or aggregations occur in certain parts of space (Fig. 2c-d), (c) *Oscillatory Patterns*, where the system never reaches a steady state so spatial patterns are in perpetual flux (Fig. 2a-b).

By a result in Potts & Lewis (2019), these parameter regimes are easily determined by calculating the eigenvalues of a matrix *A*, which we call the *pattern formation matrix*. This matrix has diagonal entries *A*_*ii*_ = −1 (for *i* = 1, …, *M*) and the entry in the *i*-th row and *j*-th column is 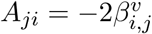 for *i* ≠ *j*. If the real parts of the eigenvalues of *A* are all negative then we are in the *No Pattern* parameter regime. If there is an eigenvalue whose real part is positive and the eigenvalue with the largest real part (a.k.a. the *dominant eigenvalue*) is a real number, then this is the *Stationary Patterns* regime. Otherwise, we are in the *Oscillatory Patterns* regime, where the dominant eigenvalue is non-real. These eigenvalues can be calculated in most computer packages, so there is no need for specialist mathematical knowledge. For example, the `R` programming language has a function `eigen()` designed for this purpose. Step-by-step instructions for the whole procedure of determining pattern formation properties are given in Supplementary Appendix D.

In Schlägel *et al.* (2019), 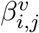 -values were inferred using SSA in all cases where *i* and *j* were of different sex, for eight different replicates (see Fig. 4 in their paper). Here, we use the published best-fit values to construct the pattern formation matrix, *A*, for each of the eight replicates. We use this to categorise each replicate by its pattern formation properties (No Patterns, Stationary Patterns, Oscillatory Patterns).

## 3 Results

### 3.1 Simulated data

When tested against simulated trajectories from diffusion-taxis equations, SSA was generally reliable at returning the parameter values used in the simulations (Fig. 1). For the Fixed Resource Model, there was just one parameter, 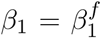 (the superscript denoting the Fixed Resource Model).

All but one of the real values lay within the corresponding 95% confidence intervals of the SSA-inferred values (Fig. 1b). The one that did not 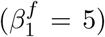 was only slightly out, so this may have been simply due to random fluctuations. SSA tended to slightly overestimate the value of 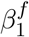 with this resource layer, particularly for higher 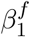 values. However, since the difference between the inferred value of 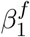 and the actual value is never very large, and within the margin of error for each individual value of 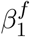, this suggests the approximations inherent in the derivation of Equation (5) from Equation (1) are acceptable for practical purposes. Fig. 1c shows the practical outcome of the small-*τ* requirement, whereby the inference over-estimates 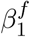 as *τ* increases. Notice also that, if *τ* is too small, the inference has large error bars, owing to minimal change in resources over the spatial extent the animal travels in time *τ*, making it hard for the SSA procedure to return a precise signal.

The SSA inference performed on the Home Range Model returned ***β***-values whose 95% confidence intervals contained the real values in > 90% of cases. Those cases where the real values lay outside the confidence intervals were always only marginally outside (Fig. 1e,f; Supplementary Fig. SF1). However, as with the Fixed Resource Model, there is a tendency for SSA to slightly overestimate the real values of 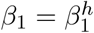 (superscript *h* for Home Range Model). The estimation of 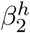 tends to be quite close to the real value unless 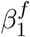 is rather large, at which point SSA starts to over-estimate 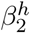 very slightly yet consistently (Supplementary Fig. SF1).

For the Home Range Model, it is interesting to examine the long-term utilisation distribution of the animal’s probability distribution, i.e. its home range. A steady-state distribution for Equation (5) is given by

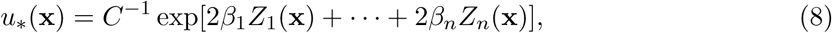

where *C* = *∫*_Ω_ exp[2*β*_1_*Z*_1_(**x**, *t*) + · · · + 2*β*_*n*_*Z*_*n*_(**x**, *t*)]dΩ is a normalising constant ensuring *u*_*_(**x**) integrates to 1, so is a probability density function. That Equation (8) is a steady-state of Equation (5) can be shown by placing *u*(**x**, *t*) = *u*_*_(**x**) into the right-hand side of Equation (5) and showing it vanishes. Note the factor of 2 before all the *β*_*i*_ in Equation (8), a phenomenon that occurred for the same reasons in a 1D version of Equation (8) in Moorcroft & Barnett (2008), where they comment on the mathematical and biological reasons behind this. Fig. 1d gives the result of plotting Equation (8) for the Home Range model with parameter values 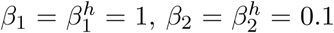. This shows how empirically-parametrised diffusion-taxis models can be used to predict home range size and shape.

### 3.2 Bank vole data

Table 1 shows the best-fit 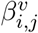-values inferred by Schlägel *et al.* (2019), together with the resulting dominant eigenvalues of the pattern formation matrix. Of the eight replicates, two of them were in the region where no patterns form, six where there are stationary patterns, but none where we predict oscillatory patterns.

**Table 1.**
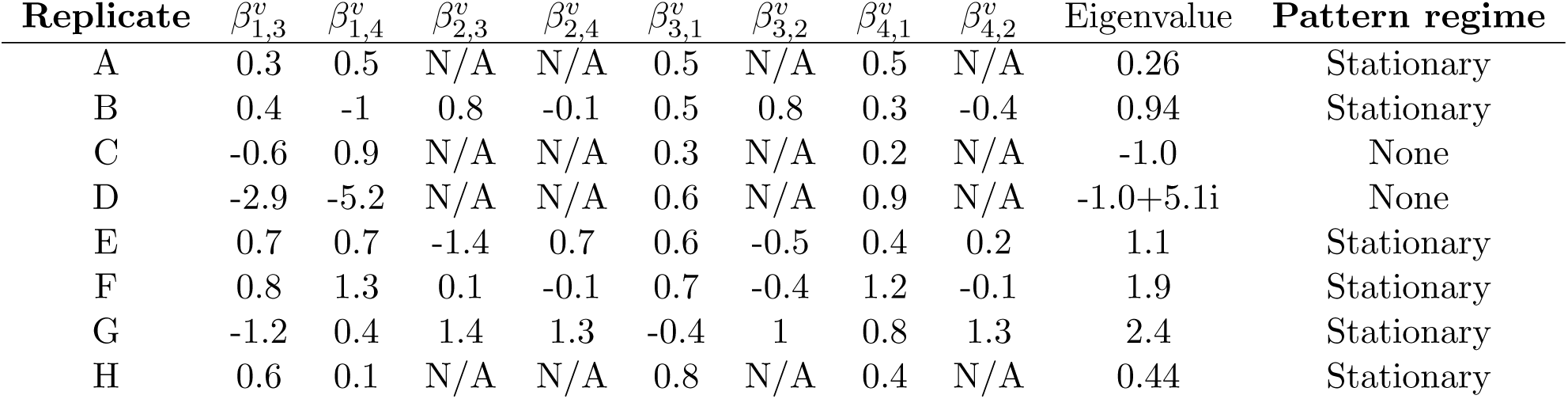
Pattern formation in bank vole populations. The first column labels the eight replicates A-H, following Schlägel *et al.* (2019). The next eight columns give the 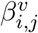-values (as defined for Equation 6) which are the best-fit values from Schlägel *et al.* (2019, Fig. 4). The penultimate column gives the dominant eigenvalue of the linearised system and the final column gives the patterning regime predicted by linear pattern formation analysis of the system of Equations (6).

Here, Individuals 1 and 2 are female, whilst 3 and 4 are male. A positive number for 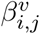 means that Individual *i* tends to move towards *j* (more precisely, *i* moves up the gradient of the utilisation distribution of *j*). For example, in Replicate A, the sole female has a tendency to move towards both males and this attraction is reciprocated. Our mathematical analysis suggests that the steady-state utilisation distribution will likely be non-uniform. One would expect, given the mutual attraction, that this would result in an aggregation of all three individuals in Replicate A. In Figs. 3a,b, we confirm this by numerically solving the diffusion-taxis equations from Equation (7) with the parameter values from the first row of Table 1 in a simple 1D domain. Note that the width of the aggregations is dependent upon the size of the spatial averaging kernel, *B*(*x*), and the exact positions of the aggregations are dependent on initial conditions (Potts & Lewis, 2019, Fig. 5). Despite this existence of multiple steady-state solutions, the general aggregation or segregation properties of the system appear to be independent of initial condition. This is proved for a simple *N* = 2 case in Potts & Lewis (2019, Sec. 4.1) and numerical evidence given for situations away from that case.

**Fig. 3.**
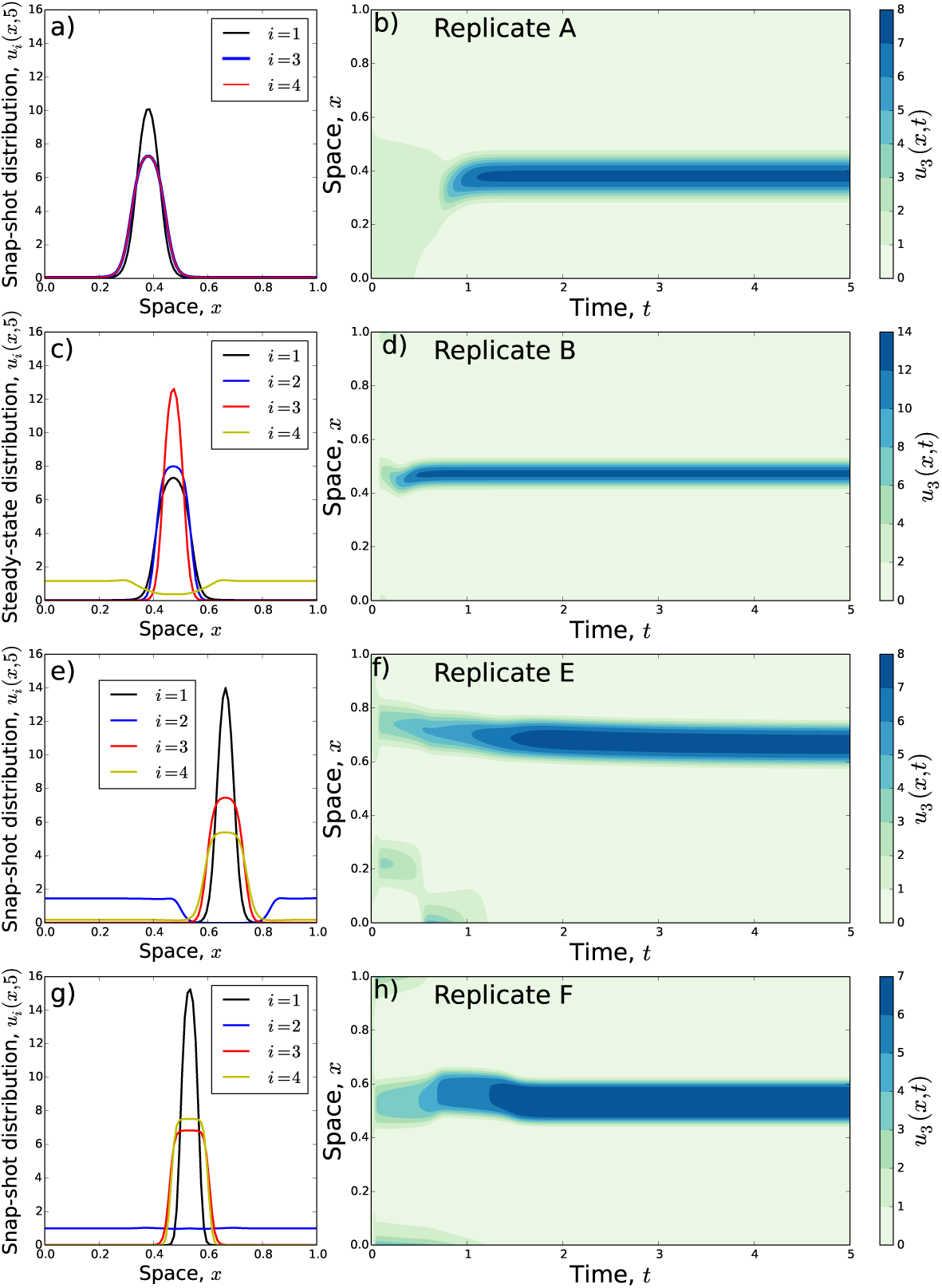
Predictions of pattern formation properties of vole replicates. These plots demonstrate whether the patterns predicted by linear analysis correspond to aggregation and/or segregation between the constituent individuals (indexed with the letter *i*). Panels (a-b) correspond to Replicate A from Schlägel *et al.* (2019), (c-d) correspond to Replicate B, (e-f) to Replicate E, and (g-h) to Replicate F. Left-hand panels give the steady-state of the distribution after solving each diffusion-taxis system numerically, with initial conditions being a small random perturbation of the homogeneous steady state (*u*_*i*_(*x*) = 1 for all *i, x*). These display the aggregation/segregation properties of the system. The right-hand panels give Individual 3’s simulated probability distribution as it changes over time.

In Replicates B, E, and F, stationary patterns are predicted to form, but the attract-and-avoid dynamics are rather more complicated, making prediction of the aggregation or segregation properties difficult to predict simply by eye-balling the 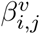-values. Numerical analysis shows that Individuals 1, 2, and 3 (both females and one male) in Replicate B tend to occupy approximately the same part of space, but that Individual 4 (the other male) tends to use the other parts of space (Fig. 3c,d).

In Replicate E, the attract/avoid dynamics given in Schlägel *et al.* (2019) show three mutually attractive parings: (1,3), (1,4), (2,4) (Table 1). This, by itself, would suggest aggregation of all four individuals. However, we also see that Individuals 2 and 3 are mutually *avoiding*, so it is not immediately obvious what the space use patterns should look like. We therefore require a numerical solution of the diffusion-taxis equations, as given in Fig. 3e,f. This reveals a three-way aggregation of both males (Individuals 3 and 4) and one female (Individual 1). The remaining female (Individual 2) strongly avoids the other three individuals, sticking to parts of space that are hardly ever used by 1, 3, and 4.

Replicate F likewise reveals complicated relationships between the four individuals. Here, numerical analysis of the corresponding diffusion-taxis system reveals an aggregation of both males (Individuals 3 and 4) and one female (Individual 1), similar to Replicate E. This time, however, Individual 2 (female) uses all parts of space, with very little tendency to avoid the others.

Replicates G and H are similar in nature to E and A, respectively. Like E, Replicate G has three mutually-attractive pairings, (1,4), (2,3), and (3,4), and one mutually avoiding pairing, (1,3). The corresponding spatial patterns (not shown) reveal aggregation of Individuals 2, 3, and 4, with Individual 1 using other parts of space. Replicate H has mutual attraction between all three individuals and, as such, leads to space use patterns (not shown) of mutual aggregation between the three individuals.

Finally, it is worth stressing that the diagrams in Fig. 3 are only there to demonstrate qualitative features of space use that diffusion-taxis analysis predicts will emerge. Principally, these are to understand whether the spatial patterns that emerge are of segregation or aggregation. However, these diagrams are not meant to represent accurate predictions of spatial patterns. Accurate predictions of space use would require incorporating into the model all the relevant resource distributions and environmental features (e.g. those in Section 2.2), together with empirically realistic initial conditions and spatial averaging kernel, in addition to details of between-individual interactions.

## 4 Discussion

We have demonstrated how diffusion-taxis equations can be parametrised from animal movement data, using the well-used and user-friendly technique of step selection analysis. The utility of such models is evidenced through two examples: (I) constructing the steady-state utilisation distribution (UD), thus relating the underlying movement to the long-term spatial distribution of a population, and (II) examining whether spatial patterns in the utilisation distribution will form spontaneously and whether these will be stable or in perpetual flux.

Despite relying on the mathematical theory of PDEs, both examples can be used without any specialist mathematical knowledge. The formula for the UD is given in a simple closed form (Equation 8), so practitioners simply need to perform SSA on their path, then plug the resulting *β*_*i*_-values into Equation (8) to infer the UD. This builds on a 1-dimensional result from Moorcroft & Barnett (2008) by generalising it to higher dimensions and linking it explicitly to the functional form given by the output of SSA. The classification of spatial distributions into ‘No Patterns’, ‘Stationary Patterns’, and ‘Oscillatory Patterns’ is done by (a) placing the *β*_*i,j*_-values into the matrix *A*, described in Section 2.3, then (b) calculating the eigenvalues, for example using the `eigen()` package in `R`. This can all be done without the need to perform technical mathematical calculations.

Our results linking the output of step selection analysis to the steady state utilisation distribution (Equation 8) are of direct application to mechanistic home range analysis (Lewis & Moorcroft, 2006). Traditionally, these were fitted to data by numerically solving a system of PDEs for a range of parameter values and searching for the best fit: a time-consuming process that requires technical knowledge of numerical PDEs. Our method, in contrast, simply requires the requisite knowledge to perform conditional logistic regression, which is both relatively quick and well-known in the ecological community.

The result of Equation (8) also makes a simple, formal link between the step selection function (SSF) and the UD that emerges from the SSF, which has an exponential form, similar to a resource selection function (RSF). This question of the UDs emerging from an SSF was examined using individual-based simulations by Signer *et al.* (2017), but our work makes this connection analytic in the case where the selection only depends on the end of the step and the turning angle distribution is uniform. Previous attempts to make this connection have started with an exponential form for the SSF and derived a rather more complicated equation for the UD (Barnett & Moorcroft, 2008; Potts *et al.*, 2014). A more recent attempt works the other way around: beginning with an exponential formulation for the UD, then deriving a movement kernel that gives the UD in the appropriate long-term limit (Michelot *et al.*, 2018). However, the resulting movement kernel does not appear in an exponential form like Equation (1). Our approach, although it relies on limiting approximation, has both a movement kernel (Equation 1) and a utilisation distribution in a similar, exponential form (Equation 8). In some sense, this is just a trivial extension of the 1D result of Moorcroft & Barnett (2008), but a useful one that has not been made explicit in the literature.

Since the predicted UD from Equation (8) is in an exponential form, similar to an RSF, it is quite straightforward for practitioners to estimate the error in this prediction and gain useful biological information about drivers of space-use patterns. First, one would subsample the data to give relocations that can be reasonably considered as independent. Then, one can re-parametrise Equation (8) using resource selection analysis on these relocation data. The *β*_*i*_-values from this re-parametrisation can then be compared with those from the SSA-PDE procedure described here.

Our results related to spontaneous pattern formation (Example II) are of particular importance with regards to species distribution modelling. These results build upon the studies of Potts & Lewis (2019) and Schlägel *et al.* (2019). The former study demonstrates the wide variety of population distribution patterns that can emerge from taxis up or down utilisation distribution gradients of other animals (including aggregation, segregation, oscillatory, and irregular patterns), whilst the latter gives a method for parametrising SSFs that describe movement responses to such gradients. The key novelty of our work with respect to the previous two is to demonstrate how the output of SSA, including from the specific SSA techniques of Schlägel *et al.* (2019), can be used to parametrise diffusion-taxis equations of the type studied in Potts & Lewis (2019). With this, we here provide the means to bridge the gap between inference on the mechanisms of fine-scale movement decisions (SSA) and predictions on resulting space-use patterns (PDEs).

Despite the wealth of theoretical work on pattern formation in animal populations over many decades [e.g. Levin (1974); Chesson (1985); Durrett & Levin (1994); Baurmann *et al.* (2007); Li *et al.* (2013)], spontaneous pattern formation is an aspect of animal space use typically ignored in species distribution models, which principally concern themselves with relating space use to environmental features. However, the literature on pattern formation gives many examples of features of spatial distributions that can arise without any need for correlation with environmental features. Perhaps part of the reason for this disparity is the perceived inaccessibility of PDEs to many ecological researchers. A major purpose of this work is to make PDEs in general, and pattern formation in particular, more widely accessible, by showing how to both parametrise and analyse PDEs using simple out-of-the-box techniques (conditional logistic regression and eigenvector calculations respectively). Of course, the analysis using such techniques is limited and much more can be done with PDEs than presented here (discussed in Supplementary Appendix F), but we hope that it will present a starting point for those who have hitherto avoided PDE formalisms.

Our use of SSA to parametrise PDEs does relies on a limiting approximation, that can affect inference. From Fig. 1c, we see that SSA tends to perform well for relatively small time-step, *τ*, but will overestimate the parameters in the PDE model as *τ* is increased. This is because the PDE moves according to the local resource gradient, merely examining the pixels adjacent to the current location. However, SSA compares the empirical ‘next location’ with a selection of control locations, which are highly likely to contain pixels that are not adjacent to the current location. This means that the movement decision may appear to be more strongly selected for than is really the case. This corroborates the idea that discretisation can lead to overestimation of selection, observed in recent theoretical work (Schlägel & Lewis, 2016b,a).

These issues of scale arise because the PDE framework in our study assumes movement along a resource gradient. One could also build a PDE model to account for attraction to resources at a distance, which is often ecologically relevant. For example, a switching Ornstein-Uhlenbeck model of resource-driven movement, such as that of Wang *et al.* (2019), has a probability distribution that evolves according to an advection-diffusion equation. It would be interesting future work to extend the framework here to incorporate such models.

## Acknowledgements

JRP thanks the School of Mathematics and Statistics at the University of Sheffield for granting him study leave which has enabled the research presented here. UES was supported by the German Research Foundation in the framework of the BioMove Research Training Group (DFG-GRK 2118/1). We thank Luca Börger for conversations and comments that have helped with this manuscript.

## Authors’ contributions

JRP conceived and designed the research. JRP performed the research, with help from UES regarding modelling the bank vole study. JRP wrote the first draft of the manuscript, and both authors contributed substantially to revisions.

## Data availability

No unpublished data were used in this study. Some results from Schlägel *et al.* (2019) were used, which can be obtained directly from Schlägel *et al.* (2019).

